# Computational Identification of Potential Inhibitors Targeting *cdk1* in Colorectal Cancer

**DOI:** 10.1101/2023.11.09.566358

**Authors:** Uchechukwu C. Ogbodo, Ojochenemi A. Enejoh, Chinelo H. Okonkwo, Pranavathiyani Gnanasekar, Pauline W. Gachanja, Shamim Osata, Halimat C. Atanda, Emmanuel A. Iwuchukwu, Ikechukwu Achilonu, Olaitan I. Awe

**Affiliations:** Department of Applied Biochemistry, Nnamdi Azikiwe University, Awka, Nigeria; Genomics and Bioinformatics Department, National Biotechnology Development Agency, Abuja, Nigeria; Department of Pharmacology and Toxicology, University of Nigeria, Nsukka, Nigeria; Department of Bioinformatics, Pondicherry University, Puducherry, India; Department of Biochemistry and Biotechnology, Pwani University, Kenya; Department of Biochemistry, University of Nairobi, Nairobi, Kenya; Biotechnology Department, Federal University of Technology, Akure, Nigeria; Protein Structure-Function Research Unit, School of Molecular and Cell Biology, Faculty of Sciences, University of Witwatersrand, Johannesburg, 2050, South Africa; Department of Computer Science, University of Ibadan, Oyo State, Nigeria; African Society for Bioinformatics and Computational Biology, Cape Town, South Africa

**Keywords:** Colorectal cancer, Cyclin-dependent kinase 1, inhibitors, natural compounds, molecular dynamics simulation

## Abstract

Despite improved treatment options, colorectal cancer (CRC) remains a huge public health concern with a significant impact on affected individuals. Cell cycle dysregulation and overexpression of certain regulators and checkpoint activators are important recurring events in the progression of cancer. Cyclin-dependent kinase 1 (CDK1), a key regulator of the cell cycle component central to the uncontrolled proliferation of malignant cells, has been reportedly implicated in CRC. This study aimed to identify CDK1 inhibitors with potential for clinical drug research in CRC. Ten thousand (10,000) naturally occurring compounds were evaluated for their inhibitory efficacies against CDK1 through molecular docking studies. The stability of the lead compounds in complex with CDK1 was evaluated using molecular dynamics simulation for one thousand (1,000) nanoseconds. The top-scoring candidates’ ADME characteristics and drug-likeness were profiled using SwissADME. Four hit compounds namely spiraeoside, robinetin, 6-hydroxyluteolin, and quercetagetin were identified from molecular docking analysis to possess the least binding scores. Molecular dynamics simulation revealed that robinetin and 6-hydroxyluteolin complexes were stable within the binding pocket of the CDK1 protein. The findings from this study provide insight into novel candidates with specific inhibitory CDK1 activities that can be further investigated through animal testing, clinical trials, and drug development research for CRC treatment.

## 1.0 Introduction

Colorectal cancer (CRC) is the third most common cancer next to lung and breast cancers with an estimated diagnosis of 2 million individuals and a mortality of approximately 900,000 annually (1, It is thought to be a disease in high-income countries; however, studies have shown the incidence of CRC to increase at a projected rate of 70% by 2030 for low and middle-income countries (3) with a 5% lifetime average-risk incidence globally (4). These gory statistics have significant implications for the public health systems both in the country and globally hence the need for a more dynamic scientific approach to the CRC burden.

The onset of CRC follows a pattern of abnormal growth of the epithelial cells of the colon and/or rectum leading to a tumor mass that fills the intestinal lumen, metastasizing into nearby organs of the abdominal area, lymph nodes, and other parts of the body (5). It usually begins as a small, adenomatous mass of cells known as polyps that form on the inside of the colon, which may become malignant or remain benign (6). As with many cancers, the cause of CRC is not known (7), however, risk factors such as age, lifestyle habits (alcohol and tobacco use), high-fat diet, predisposing medical conditions such as type 2 diabetes, and a family history of CRC have been associated with the disease course. Although traditional chemotherapy options have yielded remarkable results in CRC treatment with possibilities of relapses, limited molecular understanding of the oncogenetic initiation and interplay of biochemical mechanisms/factors have impaired the development of novel pharmacologically therapeutic anti-cancer drug agents.

Cyclin-dependent kinase 1 (CDK1) enzyme has been implicated in the oncogenesis of CRC (8). It is a serine/threonine kinase known to be the catalytic subunit of the M-phase promoting factor (MPF) complex (9) and belongs to a family of cyclic cell regulatory proteins involved in maintaining cell cycle efficiency (8, 10). Though involved in many cellular events that ensure cell survival, CDK1 specifically acts by regulating the G_2_/M and G_1_/S phases oscillating in activity between the M transition stages to bring about replication, activating checkpoints, and more recently has been essential to replication fork maintenance (11–13). Prior to initiating cell replication, CDK1 is activated by binding to cyclin B_1_ to form CDK1-cyclin B complex (fig. 1) (14). However, the over expression of CDK1 has been linked to unrestricted cell proliferation. Its over expression has been linked to unrestricted cell proliferation, tumor development, and dysregulation associated with the initiation of many cancers including colorectal cancer (8, 10, 15). Gene expression has been associated with biomarkers in complex diseases like prostate cancer (16), epilepsy (17), malaria/CoVID-19 (18) and SARS-CoV-2 infection (19). Several studies suggest that overexpression of CDK1, which is the hallmark event in many malignancies is associated with 5=fluorouracil resistance through altered activation of Wnt/β-catechin signaling, mTOR, and p53 pathways leading to poor prognosis in CRC (8, 20–22).

**Figure 1:**
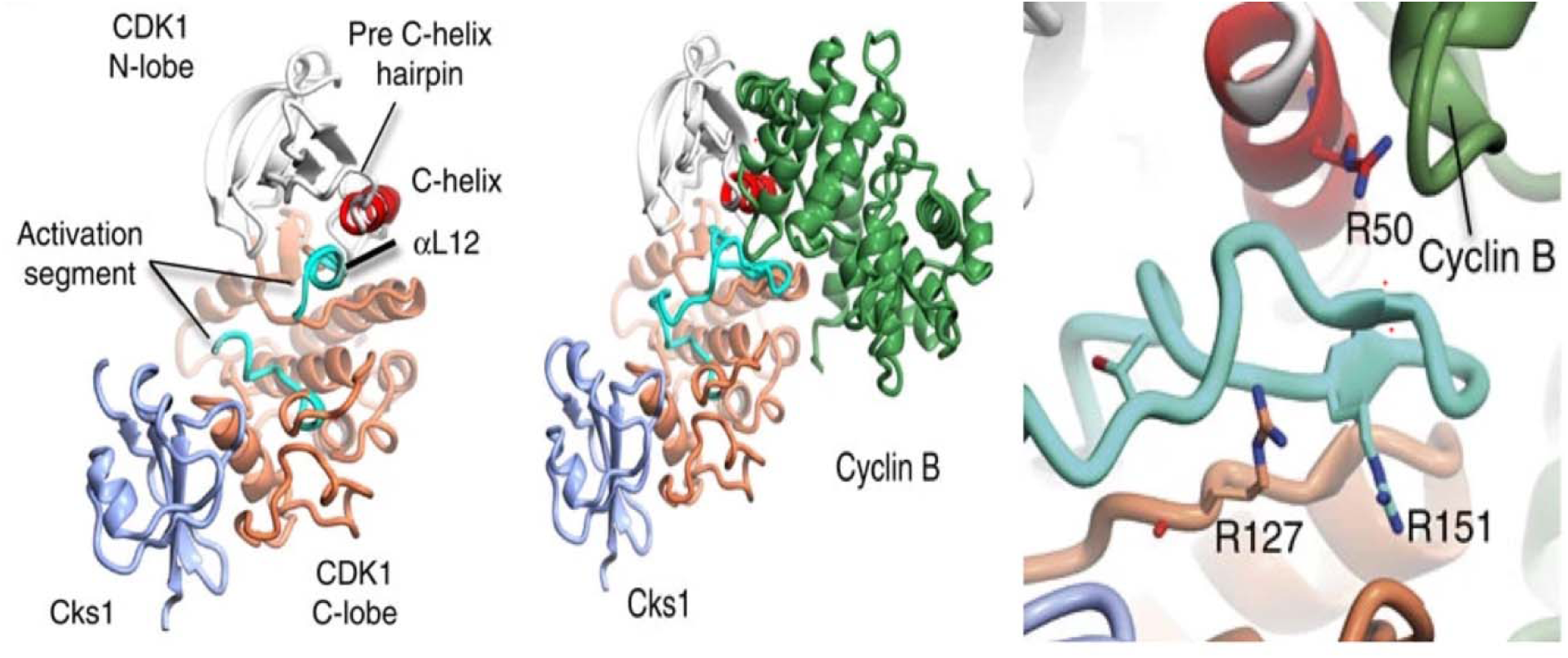
Structure of the CDKI binding to CKs1and complexation with cyclin B The complexes shown above depict (a) CDK1-Cks1 (b) CDK1-cyciln B-Cks1 (c) residues of CDK1. In each image, the following colours apply; CDKI N-lobe (white), CDK1 C-lobe (coral), Cks subunit (ice-blue), C-helix (red), activation segment (cyan), cyclin subunit (lawn green) (Brown *et al*., 2015).

Natural products have drawn a lot of attention recently as potential sources of treatments for a variety of illnesses. In recent years, bioactive compounds from plants have been proven to have opportunities in the pharmaceutical industry for therapeutic purposes (23). Resveratrol, a polyphenol has been documented for its ability to reduce or inhibit cell proliferation, reduce apoptosis and suppress angiogenesis and metastasis in cancer cells (24, 25). Also, flavopiridol, a flavonoid based substance has undergone clinical trials for its anti-cancer benefits and as a potent Cyclin-Dependent Kinase (CDK) inhibitor (26). By specifically targeting CDK1, cell cycle arrest is initiated, slowing down the growth of tumors. Compared to synthetic alternatives, natural products have fewer adverse effects.

Despite recent advances in diagnostics and treatments for CRC as well as the failure of conventional cancer treatments including chemotherapy and radiotherapy to sustain lower relapse rates, there remains a gap in treatment and increasing incidence of CRC cases. With the vast plethora of computer-aided drug design and combinatorial chemistry technologies available, a potentially wide array of therapeutic options from small molecule compounds that can target significant aspects of CDK1 activity exists. Our study investigated potential inhibitors of CDK1 *in silico* and explored their pharmacokinetic properties using a number of computational techniques.

## 2.0 Methods

### 2.1 Ligand preparation

A library of ten thousand bioactive plant compounds was compiled and downloaded from the PubChem database (https://pubchem.ncbi.nlm.nih.gov). Pubchem is a publicly available small molecule database (27, 28, 29) used in drug discovery and other applications. It contains information on over 250 million unique chemical molecules along with their descriptors. Before carrying out docking, the two-dimensional (2D) structure data file (SDF) format of the compounds downloaded from the pubchem repository were imported into Maestro (Schrodinger suite) for transformation in the LigPrep module of the software (30). Ionization states were generated at a pH range of 7±2.0 and stereoisomers were calculated at default (31).

### 2.2 Protein preparation

The three-dimensional (3D) human crystal X-ray diffraction structure of Cyclin-dependent kinase 1, (http://www.rcsb.org/pdb/home; PDB ID: 4Y72) (14) was downloaded as a pdb file and imported into Maestro suite. For appropriate structural preparation of the virtual protein using the Protein Preparation wizard of the Schrodinger software, hydrogen, bond orders, and missing side chains were added appropriately while water molecules beyond 5Å from het groups were deleted before optimization at pH 7.0+2.0. The protein structure was subsequently minimized using the optimized potentials for liquid simulations (OPLS) 2005 force field to a root-mean-square-deviation (RMSD) of 0.30Å (32).

### 2.3 Molecular docking

The compounds were subjected to molecular docking within the active site of CDK1. Using the receptor grid generation feature in the Schrodinger suite, a grid box where each ligand will be docked was generated. The binding pocket where the co-crystallized ligand is located was used to construct the defining box as well as the residue locations within the 4 Å radius of the ligand. Molecular docking was performed via the standard precision (SP) and extra precision (XP) of the Glide tool in the software (33–34). Following docking, the top compounds with optimal binding scores were selected for further studies.

### 2.4 Drug likeness and *in silico* ADME predictions

The drug-likeness of the lead compounds was evaluated by subjecting them to computational pharmacological and pharmacokinetic analyses on the SwissADME web server (http://www.swissadme.ch) (35–36), using the generated Canonical SMILES (simplified molecular-input entry system) transformations of the structures (37). Lipinski’s rule of five was used to assess the compounds based on the following parameters of molecular weight, number of hydrogen bond donors and acceptors and octanol-water partition (38–40). Pro-Tox II online server (https://tox-new.charite.de/protox_II) was used to predict the toxicity of the compounds (41).

### 2.5 Induced Fit Docking

The top four (4) ligands with the least binding scores were selected and subjected to the mixed molecular docking and dynamic protocol used by the induced-fit docking (IFD) module of the Maestro algorithm. An implicit solvent model with the OPLS 2005 force field was used to apply the standard IFD protocol to the chosen (centroid) amino acid side chains (8–10, 18–20, 31–33, 64, 80–89, 135, 145–146). The IFD procedure included a ring conformational sampling with a 2.5 kcal/mol energy barrier and a non-planar conformation penalty on amide bonds. Each ligand was given a maximum of 20 permitted postures, with the scaling for both the receptor and the ligand fixed at 0.5. The Prime Refinement technique was used to further refine residues that were within 5 Å of the docked ligand. The protein-ligand complexes with the improved structure were ranked using prime energy. For a final round of Glide docking and scoring, the receptor structures that were less than 30 kcal/mol from the minimal energy structure were submitted. In the next second docking stage, all ligands were re-docked into all revised low-energy receptor structures using Glide XP’s default settings

### 2.6 Molecular dynamics simulation

The docked complexes were further subjected to molecular dynamics (MD) simulations using the Desmond module of the Schrodinger package on a GPU-enabled Linux computer. To simulate physiological conditions, the model was created in a TIP3P (transferable intermolecular potential-three point), orthorhombic water buffered solvation box of 10 Å dimensions and 0.15M salt. The simulation model was then minimized and neutralized by adding sodium (Na^+^) and chloride (Cl^-^) ions, placed at least 20 Å from the ligands. The protein-ligand complexes were simulated for 1,000 nanoseconds in an OPLS-2005 force field. At the end of the simulation period, the root mean square deviation (RMSD), root mean square fluctuation (RMSF), radius of gyration (Rg), Ligand RMSD, solvent accessible surface area (SASA), and intramolecular hydrogen bonds (intraHB) were computed and subsequently analyzed (42–44).

### 2.7 Molecular Mechanics/Generalized Born and Surface Area Solvation (MMGBSA) Calculations

Molecular mechanics/generalized Born solvent area (MM/GBSA) calculations (45–47) were performed to estimate the relative binding energies of the selected ligands to CDK1. This method estimates the amount of free energy involved in interactions of the protein-ligand complex. The binding energy for 2000 independent snapshots of each ligand-protein complex was generated and average binding energies calculated. ΔG of each ligand was calculated following the given equation:

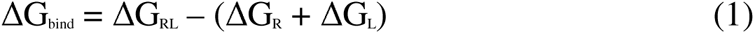

Where ΔG_RL_ represents the free energy of protein-ligand complex; ΔG_R_ is free energy of receptor, and ΔG_L_ the free energy of ligand.

The free binding energy establishes the binding affinity of ligands to a given protein. A more negative ΔG value indicates better binding of the ligand to the protein (48). The free energy of each state was calculated, using FF14SB force field, by the given equations…

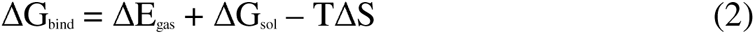

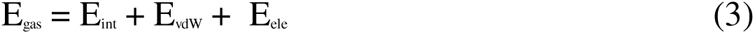

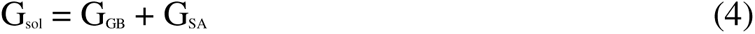

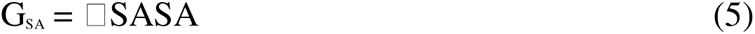

Contribution of each residue to the total binding free energy was also obtained using MM/GBSA method in Amber 18.

## 3.0 Results

### 3.1 Analysis of the binding modes of co-lig and hit compounds in CDK1 active site

We performed molecular docking study of the phyto-compounds against the CDK1 target protein. We selected the top four hits with the most favourable binding scores for subsequent studies. To validate the docking procedure, the co-crystallized ligand was re-docked into the binding pocket of CDK1. The RMSD between docked and native pose (fig. 2) is given as 0.2763. Figure 3 depicts the result from molecular docking studies of the four top-scoring compounds: Spiraeoside (-12.47 kcal/mol), Robinetin (-12.22 kcal/mol), 6-hydroxy luteolin (-12.07 kcal/mol), Quercetagetin (-12.06 kcal/mol), and the co-crystallized ligand, LZ9 (-13.99 kcal/mol). The 2D protein-ligand interaction diagrams (fig. 4) show the hydrogen and hydrophobic bond network and interactions shared between the compounds and CDK1. In this study, it was observed that all 4 hit compounds formed hydrogen bonds and hydrophobic interactions with the Ile10 residue of CDK1. Hydrogen bond interactions also existed between the compounds and CDK1 via Leu83, Ser84, and/or Gln 132 residues. Hydrophobic interactions were seen between all the compounds and Ile10, Val18, Ala31, Val64, Phe80, Phe82, Leu83, Met85, Leu135 and Ala145 residues of CDK1. LZ9 shared hydrophobic bonding with Tyr15, in addition to the aforementioned. Hydrophobic bonds observed from molecular docking were maintained for induced fit docking. Bond interactions for molecular docking, induced fit docking and molecular dynamics simulation are summarized in table 1. The complexes formed from induced-fit docking are displayed in figure 5.

**Figure 2:**
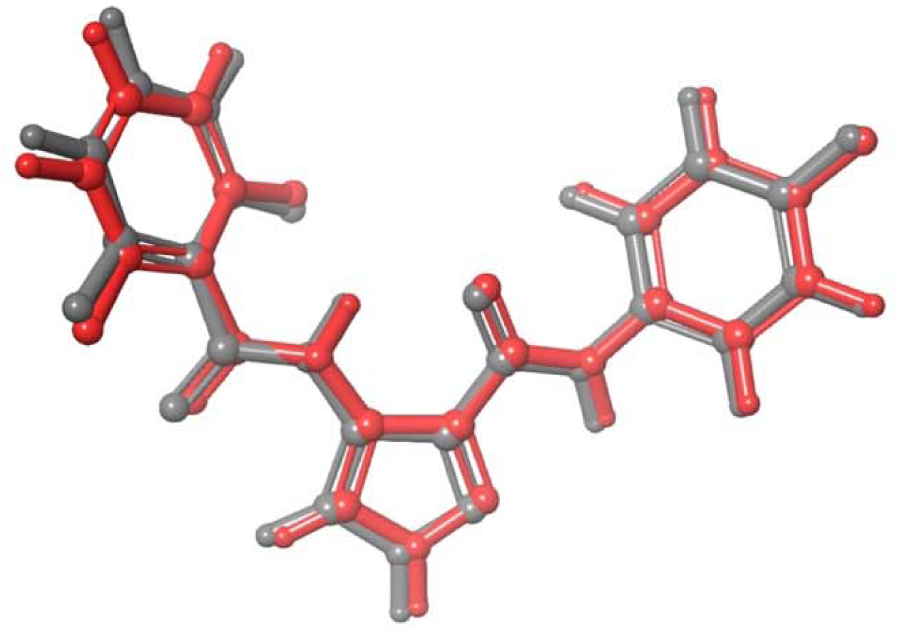
Validation of docking showing the co-crystallized ligand (in grey) and the re-docked pose (in red) with an RMSD score of 0.2763. The low RMSD value obtained demonstrates the efficacy of the docking protocol used.

**Figure 3:**
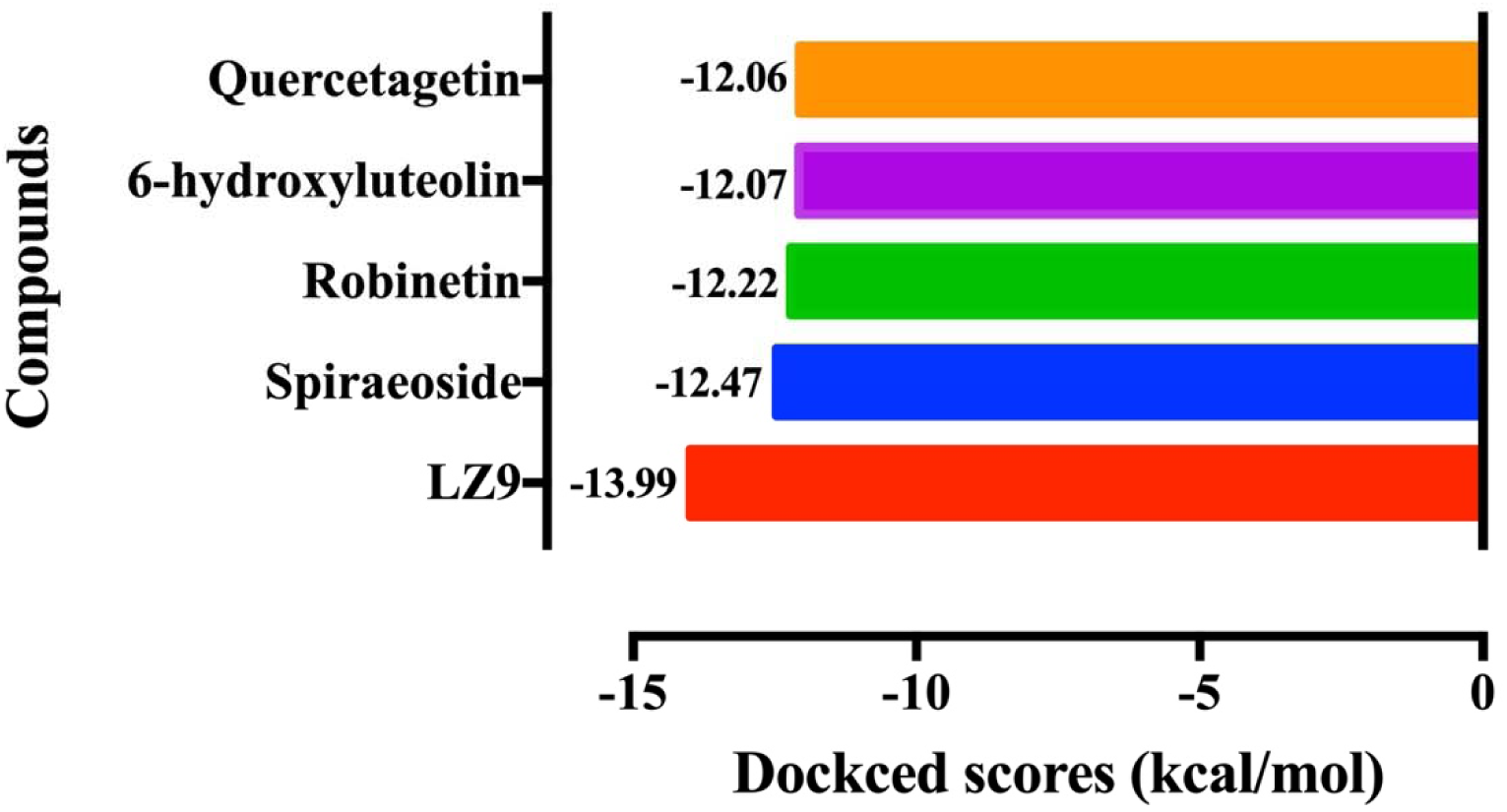
Molecular docking results of the top four (4) scoring compounds: Spiraeoside, Robinetin, 6-hydroxyluteolin, Quercetagetin and co-crystalized ligand (LZ9)

**Figure 4:**
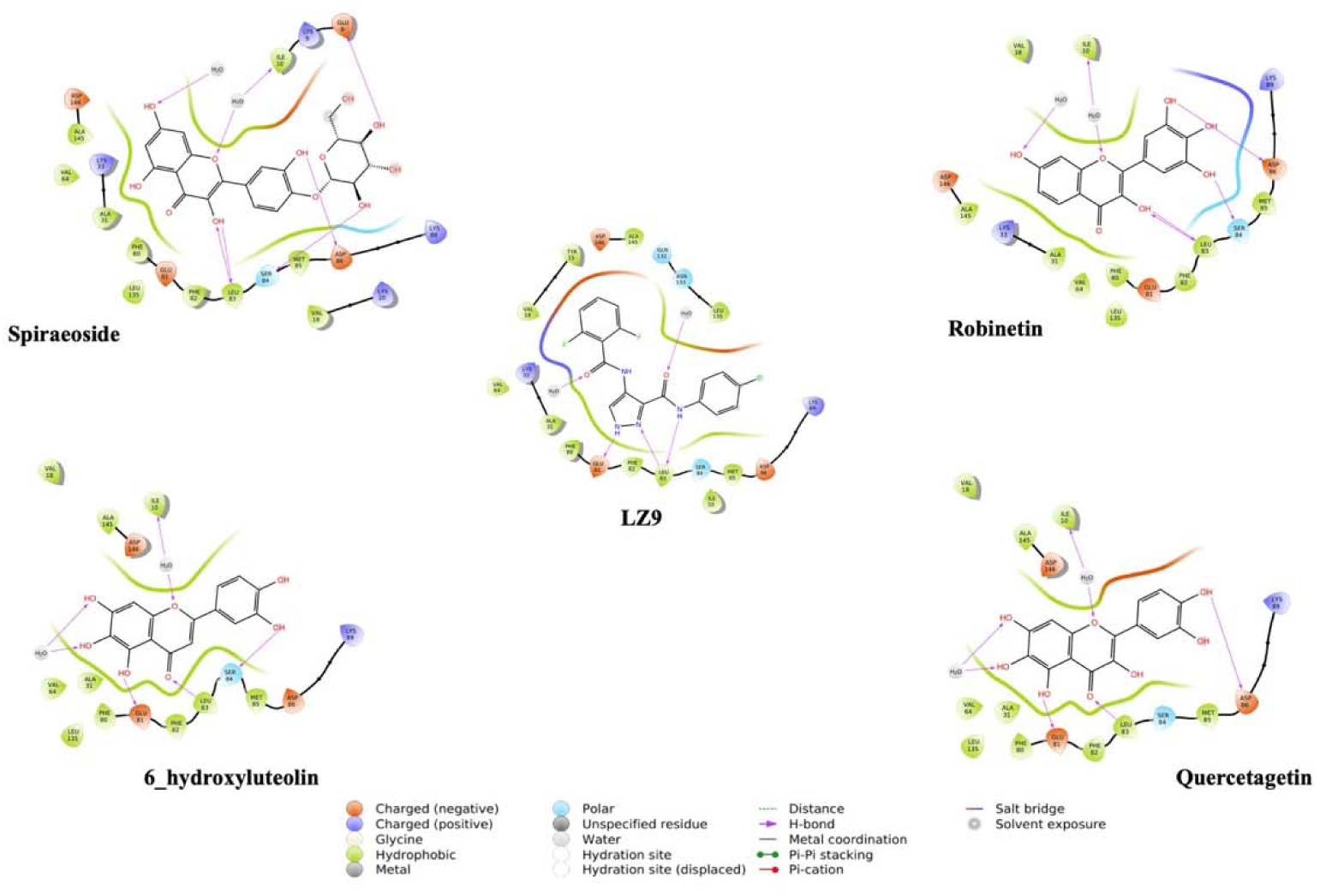
2D interaction diagrams of top scoring compounds in complex with CDK1 from molecular docking studies. Maestro 2D interaction diagram (Schrödinger) was used to generate the image.

**Figure 5:**
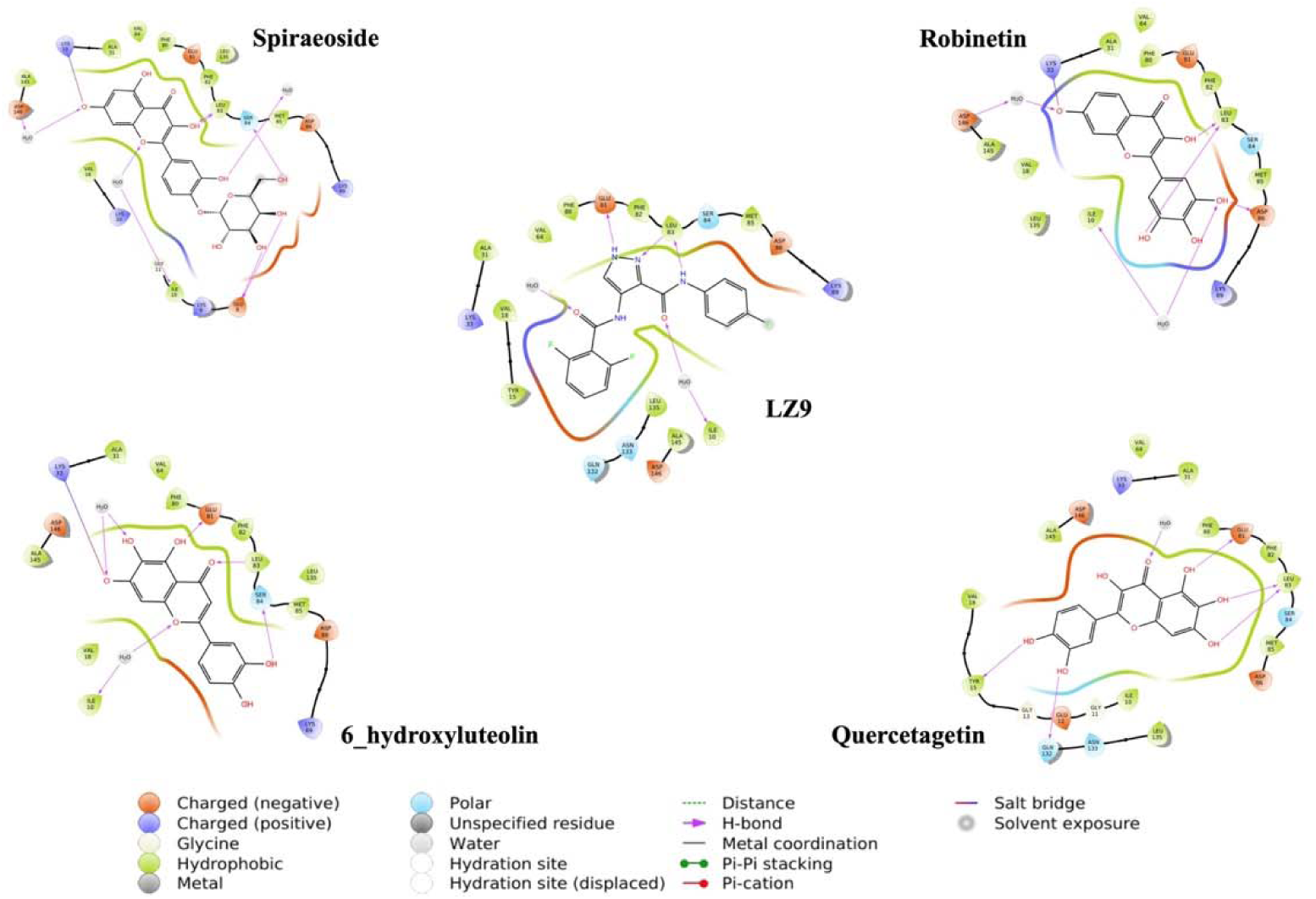
2D interaction diagrams of poses from Induced fit docking.

**Table 1:** Bond interactions between ligands and CDK1 during molecular docking, induced fit docking and molecular dynamics simulation.

| Compounds | Hydrogen bonds | | | π-π bonds |
| --- | --- | --- | --- | --- |
|  | MD | IFD | MDS |  |
| **LZ9** | Glu81, leu83 | Ile10, Glu81, Leu83 | Glu81, Leu83, Asp146 | Tyr15 |
| **Spiraeoside** | Glu8, Ile10, Leu83, Ser84, Asp86 | Glu8, Ile10, Lys33 Leu83, Ser84, Asp146 | - | - |
| **Robinetin** | Ile10, Leu83, Ser84, Asp86 | Ile10, Lys33, Leu83, Asp86, Asp146 | Thr14, Lys33, Val64, Glu81, Leu83, Arg158 | Phe80 |
| **6_hydroxyluteolin** | Ile10, Glu81, Leu83, Ser84 | Ile10, Lys33, Glu81, Leu83, Ser84 | Thr14, Lys33, Glu51, Leu83, Arg158, Phe147 | - |
| **Quercetagetin** | Ile10, Glu81, leu83, asp86 | Tyr15, Glu81, Leu83, Gln132 | Lys33, Glu51, Leu83 | - |

MD: Molecular docking; IFD: Induced Fit docking; MDS: Molecular Dynamics Simulation

### 3.2 *In silico* ADME predictions

Table 2 gives details of ADME and toxicity predictions. Physicochemical descriptors, ADME (Absorption, Distribution, Metabolism, and Excretion) parameters, and pharmacokinetic properties of the compounds, except spiraeoside, were predicted to be within the permissible parameter ranges. From toxicity prediction, spiraeoside was predicted to be inactive for all parameters. LZ9 was predicted to be hepatotoxic and carcinogenic, but not mutagenic. Robinetin, 6-hydroxyluteolin, and quercetagetin were predicted to be inactive for hepatotoxicity, while results for carcinogenicity and mutagenicity were predicted to be active. Their probability scores are shown in brackets.

**Table 2:**
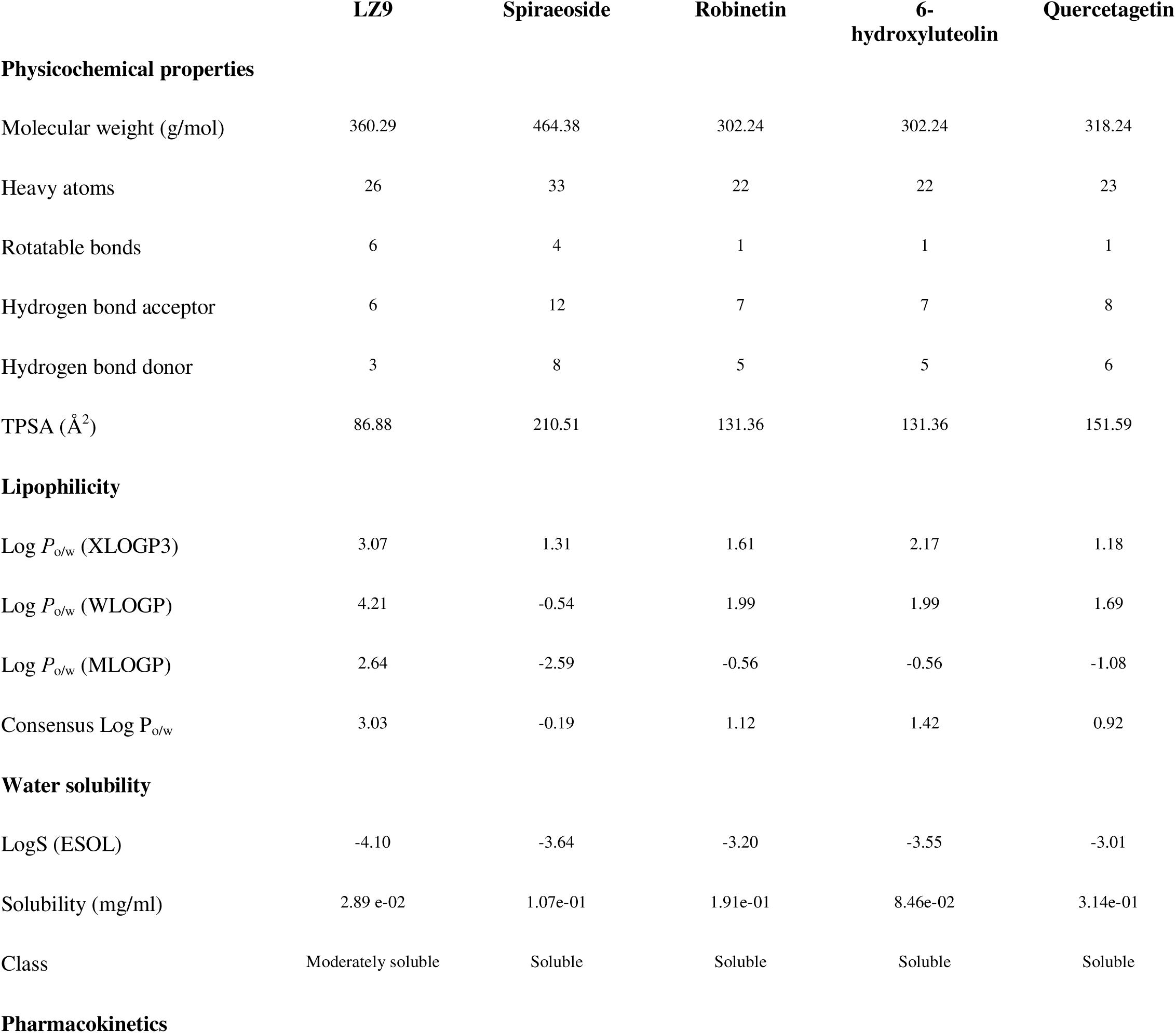
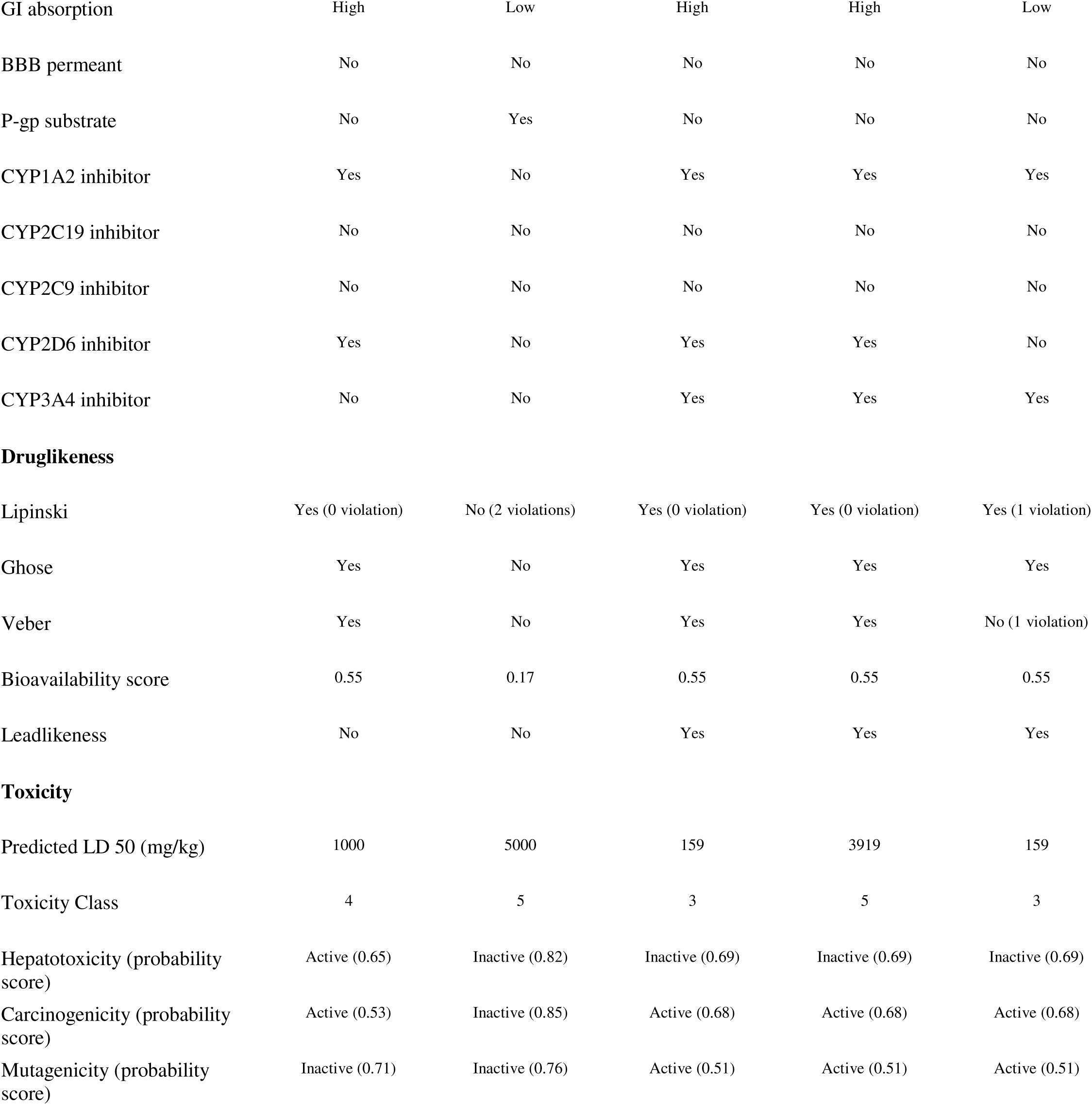
Predicted physicochemical descriptors, ADME parameters, pharmacokinetic properties, druglike nature, medicinal chemistry friendliness and toxicity of compounds.

### 3.3 RMSD analysis of CDK1 and hit compounds

The protein backbone RMSD (CCl) & ligand heavy atoms RMSD over the course of simulation time were plotted. The average values of each system were also calculated. The RMSD depicts the dynamic behavior and stability of the complexes over the simulation period and estimates the average change in atom displacement for a given frame relative to a reference frame. All protein frames were first aligned on the reference frame backbone before determining the RMSD based on the atom selection. RMSD fluctuations give insight to the stability of a ligand within the binding pocket of a protein. A gradual increase in RMSD values was observed in the first 200 nd of the app protein, after which it stabilised and had an average value of 2.48Å (fig 6A). The CDK1-LZ9 system fluctuated for the first 400 ns, after which it stabilised and had an average of 2.36Å. CDK1-robinetin stystem stabilised after 200ns with an average value of 2.40Å, while the CDK1-quercetagetin system fluctuated more, however, had an average value of 2.24Å. The most fluctuations were observed in the CDK1-6-hydroxyluteolin system which also had the highest mean value of 3.17Å. For the ligand heat atoms ((fig 6D)

### 3.4 RMSF analysis of CDK1 and hit compounds

In general, secondary structure components of a protein like alpha helices and beta strands are more rigid than the unstructured part and fluctuate less than loop regions during molecular dynamics simulation runs. Analysis of the RMSF plots (fig. 6C), show that of all the systems displayed similar patterns. Notable distinctions include quercetagetin with the highest peak value at residue 100, robinetin also displayed highest peak at residue 130, whereas LZ9 had highest peak value at residue 240. LZ9 had the least peak at residue 165 while robinetin had the least value between residues 230 and 240 where other system had highest peaks.

**Figure 6:**
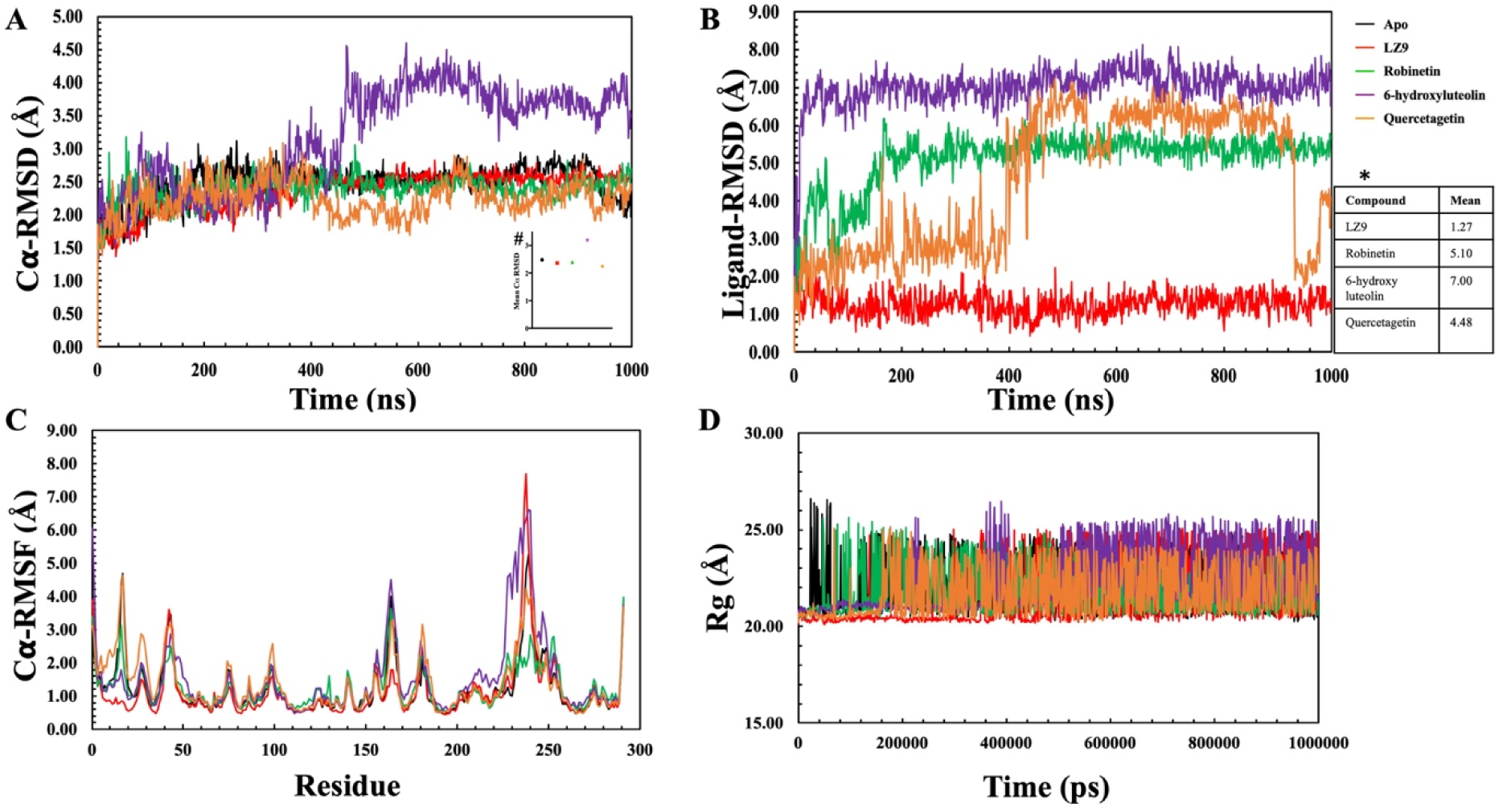
Molecular Dynamics Simulation trajectories of CDK1 in presence of ligands and absence of ligand (Apo) as a function of 1000 ns simulation time. A: Root Mean Square Deviation (RMSD) of Cl1 atoms backbone. Insert, **#** shows the mean Cl1 RMSD values. B: Root Mean Square Deviation of the ligands with respect to CDK1 protein. The plot displays the RMSD of the ligands after measuring the RMSD of the heavy atoms in the ligand and aligning the protein-ligand complex on the reference protein backbone. If the reported values are noticeably greater than the protein’s RMSD, the ligand has probably diffused away from its initial binding site. Insert, **\*** shows the mean RMSD values C: Root Mean Square Fluctuation (RMSF) per residue over time. D: Radius of gyration for the Apo state and CDK1-ligand complexes .

### 3.5 Radius of Gyration, Solvent Accessible Surface Area and Intra-hydrogen bond analysis

The radius of gyration for all systems is shown in figure 6D. When comparing the plots of the apo form to the CDK1 bound states, there is no apparent alteration to the global protein compactness in response to ligand binding. The solvent-accessible surface area describes the surface area of a molecule accessible to water. In addition to examining the impacts on the molecule, it expresses the degree to which a simulated protein interacts with solvent molecules. Evaluating the SASA value is an important way to estimate conformational changes caused by complex interactions. This is essential for characterizing protein-ligand interactions and understanding structural changes during simulations. Solvent Accessible Surface Area (SASA) plots (fig. 7A) show that LZ9 and quercetagetin have high SASA values, while robinetin and 6-hydroxyluteolin have lower SASA values. Intramolecular hydrogen bonds show the number of internal hydrogen bonds (HB) within a ligand molecule and give insight into the behavior of ligands in the active site of a protein. They can also stabilize conformations that are conducive to protein binding, thus increasing the affinity for the receptor. Intra-hydrogen bond (intraHB) plots (fig. 7B, C, D) reveal that LZ9 had intraHB interactions almost throughout the simulation time, 6-hydroxyluteolin and quercetagetin had more intraHB interactions at the beginning of the simulation, while robinetin maintained a value of zero, that is zero intraHB interactions throughout the simulation time.

**Figure 7:**
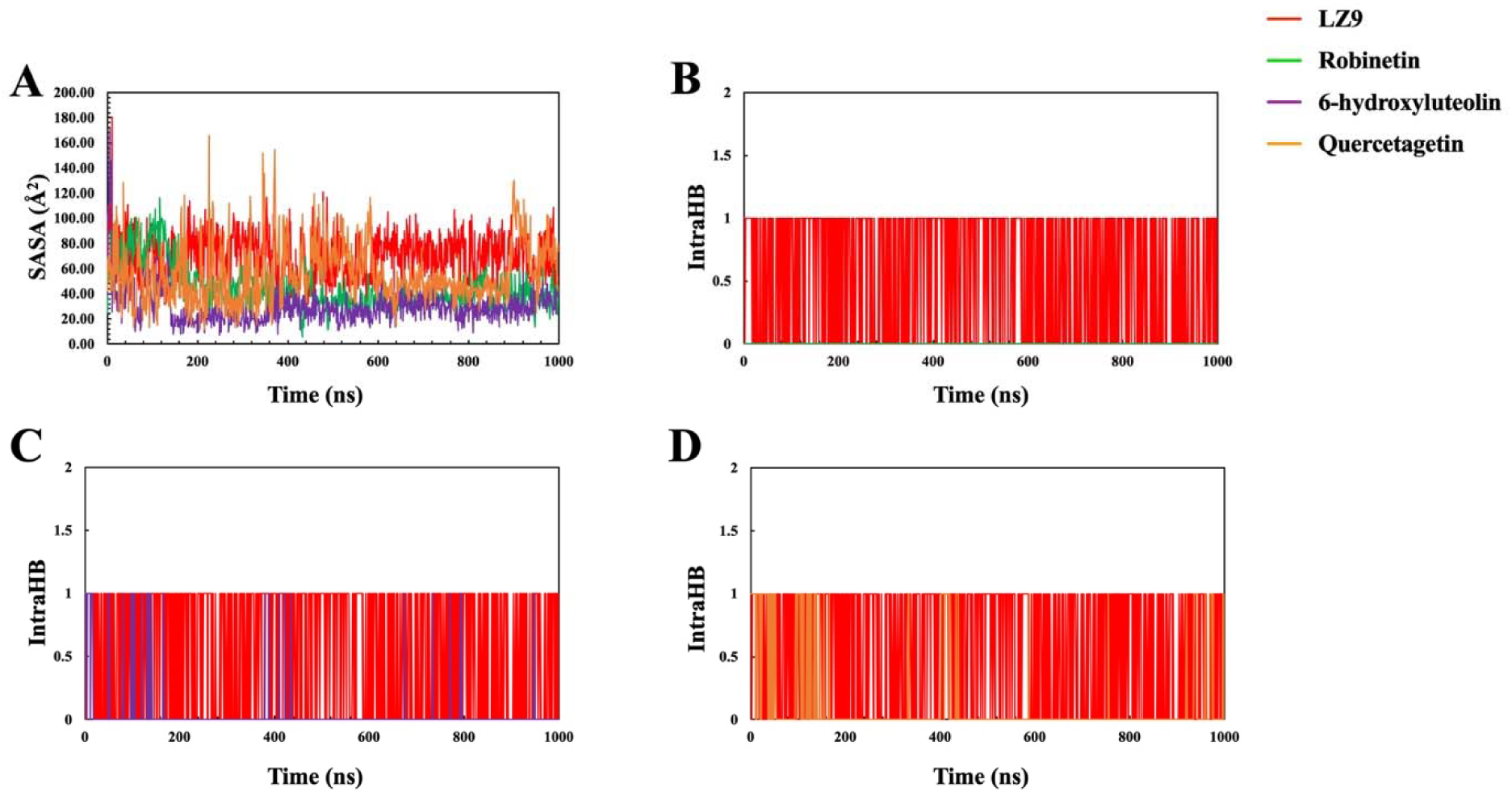
A: Solvent Accessible Surface Area (SASA) plot. Intramolecular Hydrogen Bonds (intraHB) plot showing the number of internal hydrogen bonds within the ligand molecules – Robinetin (B), 6-hydroxyluteolin (C), Quercetagetin (D), compared with LZ9

### 3.6 Protein-ligand contact

Figure 8 shows a schematic of the detailed ligand atom interactions with the protein residues. The interactions in this trajectory that occur for more than 30% of the simulation period are displayed. LZ9 and robinetin make pi-pi contacts with Tyr15 and PHE80 for more than 50% of the time. All the ligands made contact with Leu83 via hydrogen bonds; LZ9 and robinetin form hydrogen bonds with Glu81, while 6-hydroxy luteolin6-hydroxy luteolin and quercetagetin form hydrogen bonds with Glu51. Robinetin, 6-hydroxyluteolin, and quercetagetin form salt bridges with Lys33. All the ligands show relatively high solvent exposure. Protein-ligand contacts as illustrated by bar charts (fig. 9) during the simulation, show that hydrogen bonds, hydrophobic interactions, and water bridges were predominant in LZ9. The other ligands had ionic interactions in addition to the aforementioned. All compounds exhibited strong hydrogen bonding on Leu83 and hydrophobic interactions on Leu135. They also showed a water bridge on residue Asp146. Only the lead compounds showed ionic bonds at residue Lys33.

**Figure 8:**
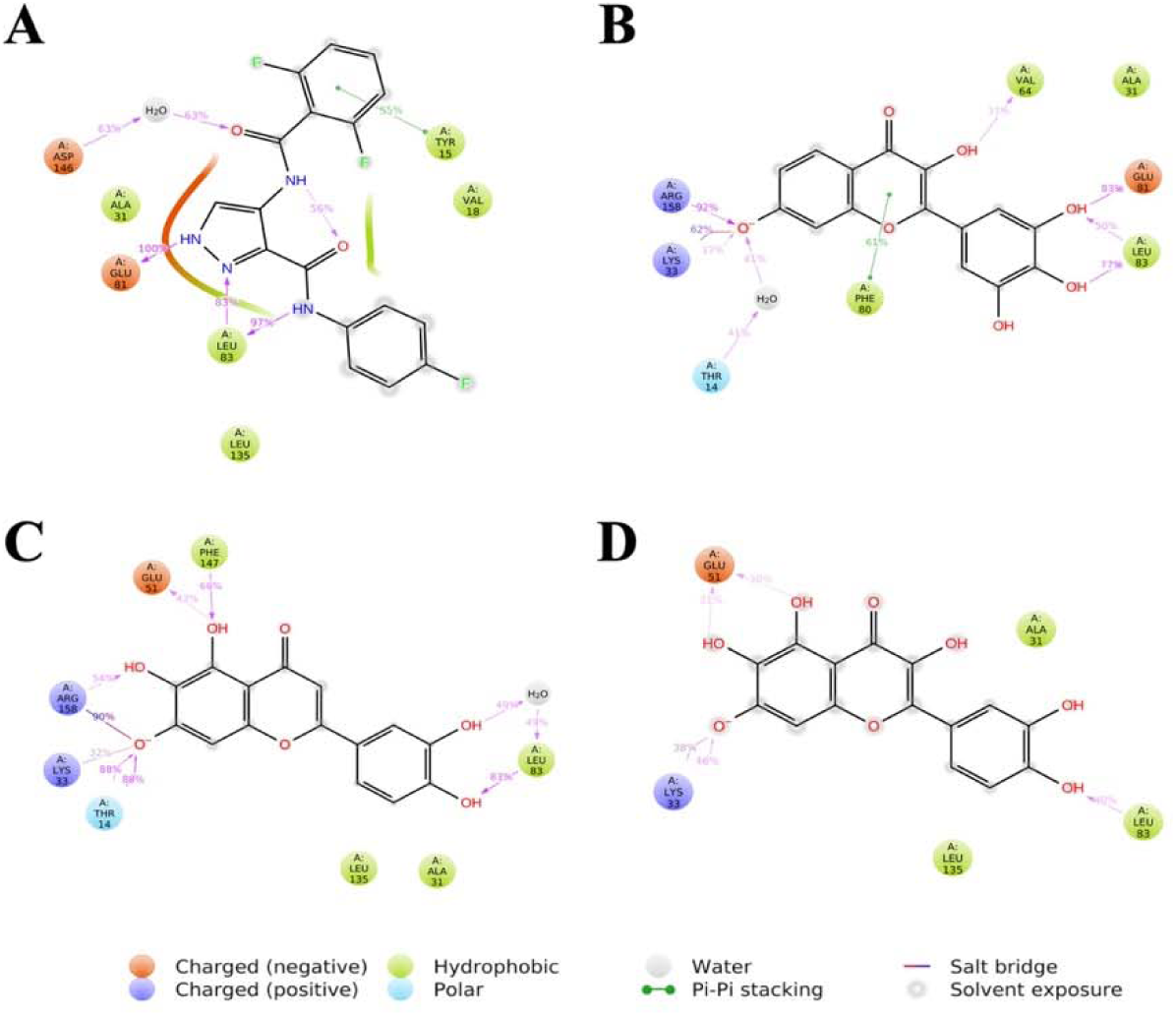
A schematic of detailed ligand atom interactions of LZ9 (A), Robinetin (B), 6-hydroxyluteolin (C) and Quercetagetin (D) with the protein residues of CDK1. Interactions that occur more than 30.0% of the simulation time in the selected trajectory (0 through 1000 ns) are shown

**Figure 9:**
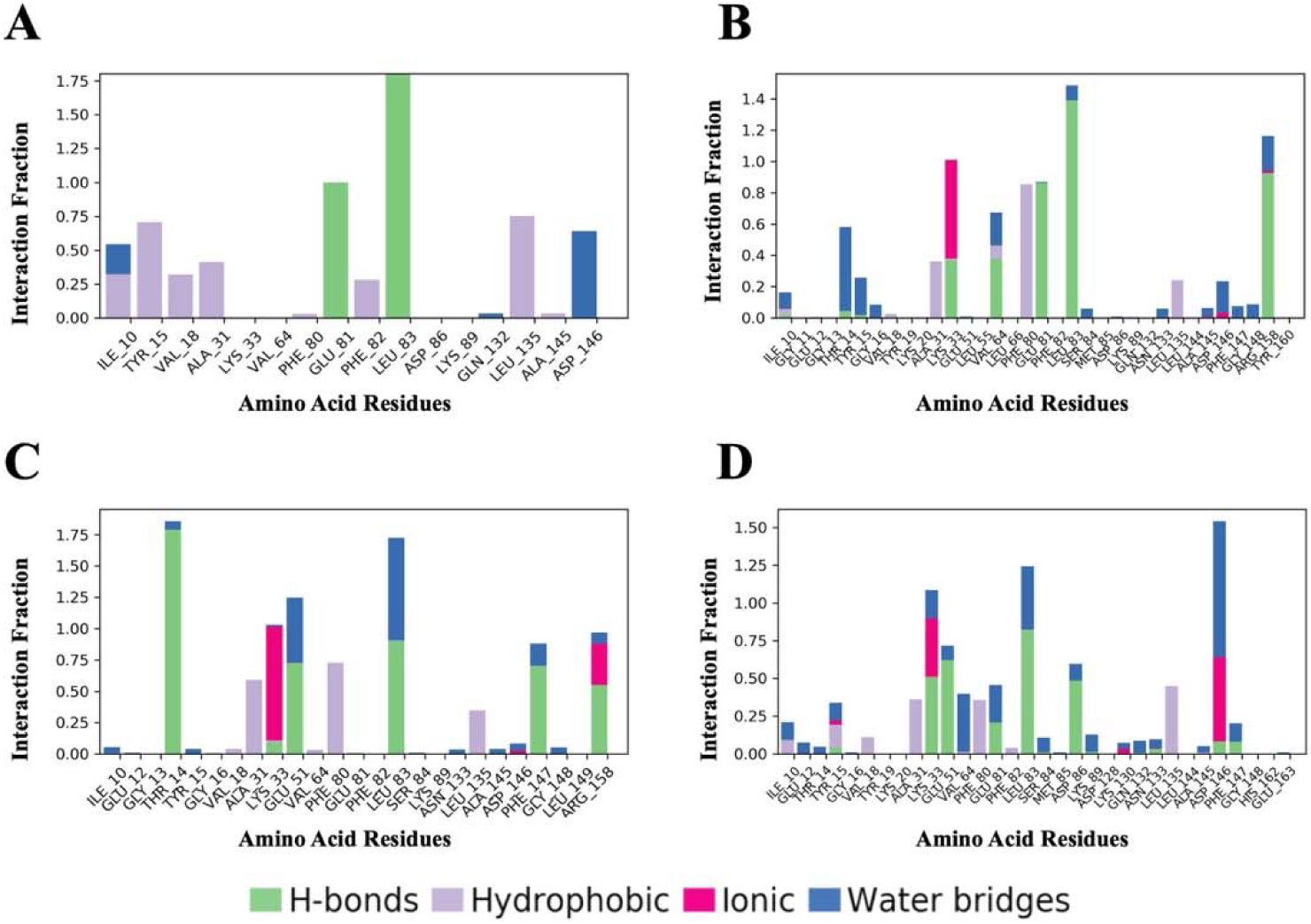
Protein-Ligand Contacts. A: LZ9. B: Robinetin. C: 6-hydroxyluteolin. D: Quercetagetin. Contact interactions can be categorized by type and summarized, as shown in the plot. Protein-ligand contacts are categorized into four types: Hydrogen bonds, Hydrophobic, Ionic and Water Bridges. The stacked bar charts are normalized over the course of the trajectory: for example, a value of 0.7 suggests that 70% of the simulation time the specific interaction is maintained. Values over 1.0 are possible as some protein residue may make multiple contacts of same subtype with the ligand.

### 3.7 Potential energy

Figure 10 shows the potential energy plot of the systems in the apo state and in complex with the ligands. When bound to LZ9, robinetin and quercetagetin, we observe energies in the same range as that of the apo state; -115,500 to -117,000 kcal/mol. On the other hand, when bound to 6-hydroxyluteolin, a lower energy state of -144,000 kcal/mol is observed.

**Figure 10:**
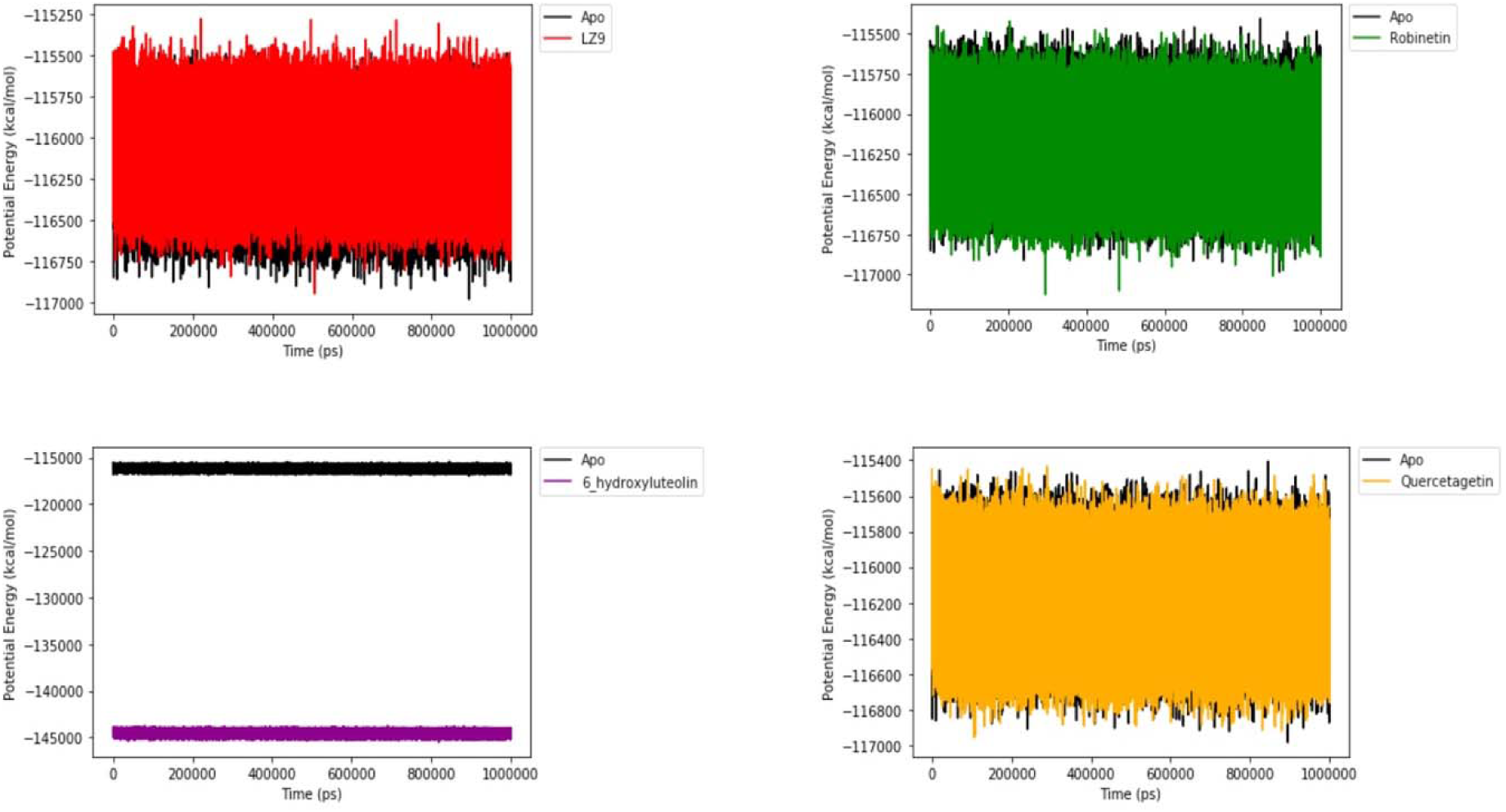
Potential energy plot of 1000 ns simulation time. The plots show the energies of the systems in apo and bound states.

### 3.8 Free binding energy

Robinetin and 6-hydroxyluteolin exhibited good ΔG values of -30.40 kcal/mol and -30.63 kcal/mol, respectively (table 3). The co-lig, LZ9, had a value of -34.96 kcal/mol. Quercetagetin had a binding energy of -18.52 kcal/mol. Energy contributions of active site residues responsible for stabilizing the CDK1-ligand complexes are shown in figure 11. Worthy of note is Glu81 as the residue showing the highest energy contribution in the CDK1-LZ9 complex, while Asp86 is the residue showing highest binding energy contribution to the CDK1-robinetin and CDK1-6-hydroxyluteolin complexes. The CDK1-quercetagetin system has Asp146 as the active site residue contributing the highest energy. Visual inspection of the figure shows that Van der Waals forces contribute more to the interaction between CDKI-LZ9 and CDK1-6-hydroxyluteolin complexes. Robinetin has more electrostatic energy contributing to the stability of the complex it forms with CDK1. Quercetagetin forms weak interaction with CDK1 as a result of the overall low Van der Waals forces and electrostatic energy.

**Table 3:**
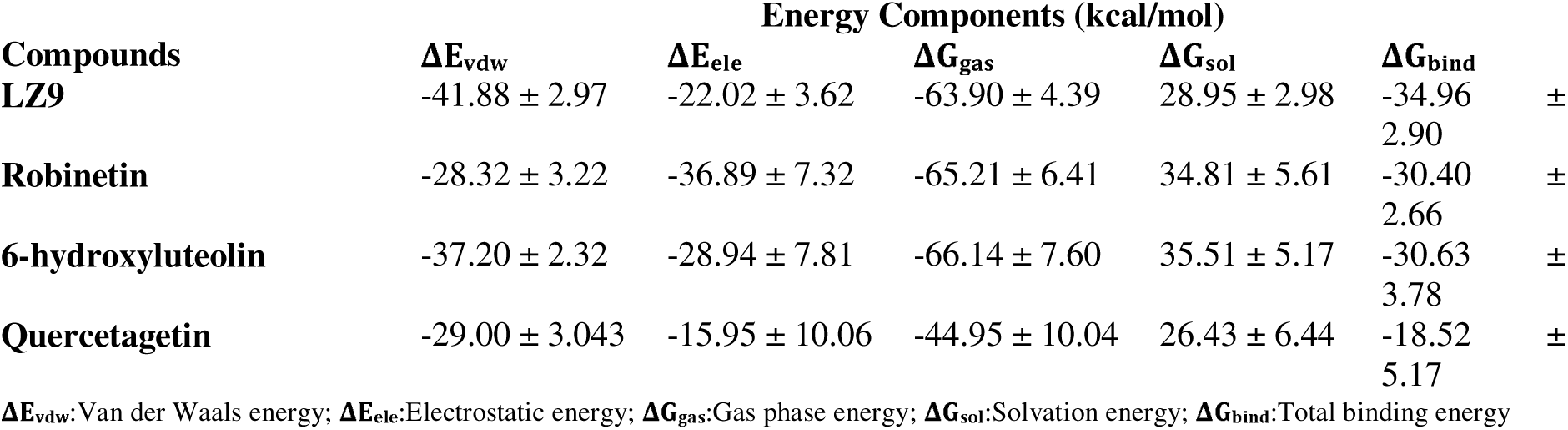
Energy component calculations obtained by MM/GBSA method.

**Figure 11:**
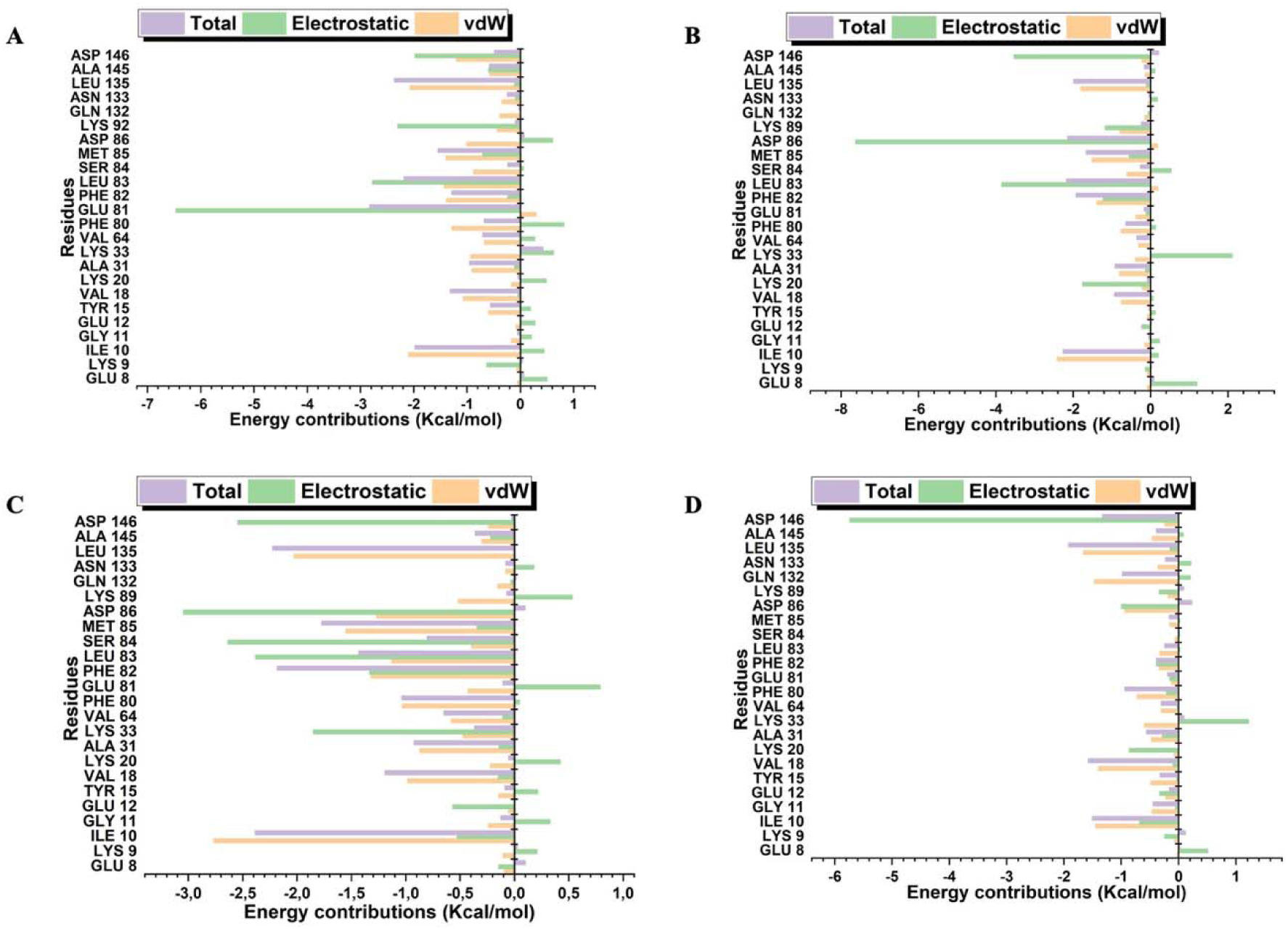
Active site residues and the resultant energy contributions in stabilizing the complexes formed between CDK1 and LZ9 (A); robinetin (B); 6_hydroxyluteolin (C); and Quercetagetin (D).

## 4.0 Discussion

The prevalence of colorectal cancer is fast becoming a public health concern (2). Development of new pharmacologically therapeutic anti-cancer drug agents has been hampered by the limited molecular understanding of the oncogenetic initiation and the interaction of biochemical mechanisms/factors, although traditional chemotherapy options have produced impressive results in the treatment of CRC with the possibility of relapses.

Natural products are a valuable source of novel therapeutic leads given that they contain a wide variety of chemical components (49–50). They have been essential in the development of new drugs, particularly for the treatment of cancerous disorders (51–52). The compounds identified from this study were found to be in the class of flavonoids. There is growing evidence that flavonoids may be useful in the prevention and treatment of cancer and they are also reported to have a variety of anti-cancer effects, including inducing apoptosis in cancer cells, inhibiting cell proliferation, and may enhance chemotherapy (53). Flavonoids have also been reported to play a potential role in colorectal cancer therapies (54).

Given the importance of cyclin-dependent kinases (CDK) in cancer, targeting CDKs is a viable strategy for combating the cancer menace (8). Because of the significant role it plays in one of the phases of cell cycle control, which is a crucial one for the growth of tumor cells, specifically targeting CDK1 became a strategic focus for optimized treatment. The drug discovery pipeline greatly benefits from the use of computational screening techniques (55). In recent years, computer-aided drug development methods have been widely used to find lead compounds with therapeutic value. These technologies include molecular docking, *in silico* pharmacokinetic profiling, and molecular dynamics simulations (MDS) (56–57). MDS are anticipated to become more significant as new pharmacological therapies are developed (58). Several studies now use these computational methods to quickly identify potential inhibitors of key proteins in disease progression as treatment options (8, 59–61) others have identified experimentally validated leads (62–63). Here, we used these computational techniques to screen phyto-compounds and identify possible inhibitors of CDK1.

Molecular docking is a computational method used to predict the preferred binding orientation of a ligand to a receptor. The software algorithm used predicts possible binding orientations and provides scores ranked according to the predicted binding affinity between the receptor-ligand complex (56,64). To better understand the binding dynamics of the four compounds selected in this study to CDK1, they were subjected to molecular dynamics simulations (MDS). Prior to running simulation studies, we carried out induced fit docking (IFD) (65). Induced fit docking is a more realistic model of how proteins and ligands interact in nature as it considers the flexibility of the protein and ligand, allowing them to move and rotate during the docking process, resulting in more accurate predictions of the binding of ligands. For traditional rigid docking, the proteins and ligands are treated as rigid bodies, and the docking algorithm attempts to find the best orientation of the ligand in the protein’s binding site. However, in reality, proteins are flexible molecules, which can undergo conformational changes upon ligand binding. Both minor backbone relaxations and substantial side-chain conformational changes in the receptor structure are accounted for by the induced-fit docking (IFD) technique (66–67).

All systems ran through the complete one thousand nanoseconds duration during MDS, except spiraeoside, which ran for only 700 ns. It could be that due to conformational changes in the protein, spiraeoside left the binding site of the protein during the simulation run. For this reason, the post-simulation analysis could not be completed, consequently, results for spiraeoside were excluded.

It is important to pay attention to the protein’s RMSD value to properly infer structural changes occurring in a protein which could affect ligand binding. CCl-RMSD demonstrated a stable structural conformation across the course of the simulation, showing that it was balanced. The low RMSD values of LZ9, robinetin and quercetagetin indicate low fluctuations and suggests that the system maintained stability during simulation and the protein did not undergo significant structural changes as a result of binding of the ligands. The higher RMSD value of 6-hydroxyluteolin system might suggest that the binding of this ligand might have perturbed the structure of the CDK1 protein.

Ligand RMSD provides insight into how stably the bound ligand sits within the protein binding pocket. The Ligand RMSD results show that LZ9, robinetin, and 6-hydroxyluteolin did not move much, therefore forming a stable complex with CDK1, unlike quercetagetin which had a lot of fluctuations. The fluctuations observed with quercetagetin after 400ns tell us that it is not tightly bound to the protein and has possibly diffused away from the binding pocket, so it forms an unstable protein-ligand complex and is less likely to perform its intended function.

RMSF measures the structural flexibility of a protein and characterizes local perturbations along a protein chain (68). It also provides information on how each amino acid residue is changing relative to its reference position during simulation (69). Here, we analysed the RMSF of the CDK1 protein in the apo & bound states. Peaks represent regions of the protein where the simulation-induced fluctuations were greatest (70). The spikes observed from RMSF results may have been caused by increased flexibility within the protein’s loop regions. It may be inferred from the RMSF result that CDK1 has a great deal of flexibility to accommodate the ligand at the binding pocket.

The radius of gyration measures how compact a protein structure is and is equivalent to its principal moment of inertia, describing the common separation of each scattered element from the molecule’s mass center (71). During simulation, a more rigid structure is represented by lower Rg values (72). Since not much difference was observed between the Apo and bound states, we can say that the binding of the ligands did not interfere with the compact structure of CDK1. In this study, the Rg, RMSD, and RMSF were correlated, and this relationship shed light on how complex compressibility affected the variations in complex RMSD and the residual fluctuations seen in RMSF. No noticeable changes are observed in the bound and unbound states of CDK1, suggesting that the protein structure retained its compact conformation. Our results indicate that the surface area of LZ9 and quercetagetin have higher exposure to water than robinetin and 6-hydroxyluteolin. This result is in tandem with that observed from the ligand-atom interaction diagram.

Hydrogen and hydrophobic bonds play key roles in protein-ligand interactions. Crucial hydrophobic and/or hydrogen bonds, including Ile10, Phe80, Leu83, and Ser84 between CDK1 and the lead compounds that were first seen after molecular docking were maintained during MDS. Phe80 has previously been described as a hydrophobic checkpoint enabling water molecules to gain access to the catalytic portion of the ATP catalytic domain for an additional hydrogen bond (73). Robinetin is seen to form a pi-pi bond with Phe80 during simulation run, which may further account for its strong inhibitory properties. 6-hydroxyluteolin also forms a bond with Thr14 for 88% of the simulation time. Inhibition of Thr14 by small molecule compounds has been proposed as a promising option for the regulation of the G2/M transition of the cell cycle (74). Inhibition of CDK1 due to the phosphorylation of the kinase on Tyr15 residue by flavonoids (75) as well as induction of G_2_/M phase arrest and apoptosis by the flavonoid tamarixetin on human leukemia cells (76) have previously been reported.

The hydrophobic and hydrogen bonds formed between the compounds and CDK1 may play a vital role in stabilizing the complexes for the compounds to exert their inhibitory efficacies (77). Ionic bonds observed on only the lead compounds might also play a significant role in their inhibition of CDK1. Interestingly, the lead compounds are all of the flavonoid class; another study by (78) also gave insight into flavonoids as CDK1 inhibitors. Natural flavonoids such as Luteolin and Daidzein have also been found to inhibit CDK1 (79).

Higher free energy values in fact correlate with more preferable binding affinities of a ligand to a protein. Robinetin, which had good binding energy to CDK1, as revealed by the dG values, might have been a result of higher electrostatic and van der Waals energy contributions. Interactions between robinetin and Asp86 residue which was observed from molecular docking and induced fit docking results was maintained after the MM/GBSA run. We can say this interaction might be a key factor to stablizing the CDK1-robinetin complex. Energetic analysis reveals that van der Waals interaction and non-polar contributions are favorable in the formation of complexes and amine group of the ligand, which plays a crucial role for binding (80–81).

The physicochemical properties of a compound are important parameters in the pipeline of drug development as they directly affect the absorption, distribution, metabolism, excretion, and toxicity profile (ADMET) of a drug candidate (82). The drug-like and ADMET predictions are used in identifying properties of a ligand suitable for formulation into drugs. Even though all the compounds are adequately hydrophilic and lipophilic, robinetin and 6-hydroxyluteolin showed high intestinal absorption. This implies that the compounds are capable of achieving pharmaceutical and therapeutic bioavailability at the intestinal linings and target cells in the colon. This is further corroborated by their bioavailability scores. A study on natural compounds as inhibitors of CDK1 by Saikat (83) also reported similar ADME properties.

Lipinski’s Rule of five is used to assess and predict the drug-likeness of compounds based on oral absorption, membrane permeability and bioavailability (38). Although, parenteral administration may be applicable in most cases of cancer treatment, oral administration is a valuable treatment option for cancer patients as it is more convenient and reduces the frequency of their clinic visits or venipuncture. In recent times, oral anti-cancer drugs such as capecitabine, topotecan and vinorelbin are successfully used in the treatment of a variety of tumours like colon, breast, ovarian or lung cancer (84). With the advent of targeted deliveries, most of the drug molecules approved as oral medications for cancer treatment including hormone therapies such as, dasatinib, sorafenib, imatinib, lapatinib and abiraterone have resulted in good clinical outcomes (85–86).

Despite the fact that natural products frequently violate these rules due to their complexity, they still serve as reliable sources for novel drug discovery and development (87). Some anti-cancer agents such as vincritine, irinotecan, paclitaxel and etoposide, widely used for decades in cancer treatments originate from natural compounds (88).

The ProTox–II server is a web-based tool that predicts the potential toxicity of a given small molecule compound on various organs and systems in the body. The toxicity is predicted using a machine learning model that has been trained on a sizable dataset of toxic and non-toxic molecules. The toxicity prediction and probability score are considered important parameters for predicting a compound’s toxicity (89). The probability score can be used to evaluate the prediction’s level of confidence. A forecast with a high probability score is more certain to occur, whereas one with a low probability value may not. Additionally, if the organ toxicity and/or toxicity end points of a compound are classified as active, it suggests that the compound may have harmful effects if it is ingested, inhaled, or otherwise exposed to the body. Conversely, "inactive" could mean that the compound has not been predicted to have a toxic effect. Early identification of toxicity in the drug development steps is critical in reducing late-stage attrition rates (90–91). The predicted toxicity class exhibited by the compounds indicates that they can be administered within the accepted safety doses. Although the test ligands are predicted to be non-hepatotoxic, lead optimization may be required to modify the other toxicity endpoints.

## 5.0 Conclusion and Next Steps

In this study, computational techniques including molecular docking, induced fit docking, molecular dynamics simulation and MM/GBSA were used to study the potential inhibitory activities of natural compounds against CDK1. The findings provided insight into molecular interactions of significant residues involved in inhibitory CDK1 activities through molecular docking and molecular dynamics simulation results. The current investigation of small molecule inhibitors in the activity of the CDK1 explored viable options for the roles of Robinetin and 6-hydroxyluteolin in the pathogenesis of CRC. This research, however, provides a basis for further examination of potential phytochemicals for medical intervention of several cancers. This study aimed to identify potential small molecule inhibitors of CDK1 hence the use of molecular modeling approaches was deployed to highlight compounds of strong binding energies, good spatial amino acid residue arrangement, and better pharmacokinetic properties among others. The MD simulations showed that the hit compounds stably interacted with the CDK1 enzyme and maintained firm positions within the binding pocket of the enzyme over the simulation timeframe. The findings of this study should undergo experimental validation to confirm the interaction between the top ligands identified in our research and the target protein. Additionally, the ADME/Tox properties that were predicted using computational models need to be verified through experimental testing.

## Availability and Requirements

Project name: e.g. In silico identification of CDK1 inhibitors in colorectal cancer

Project home page: https://github.com/omicscodeathon/cdk1_crc

Operating system(s): Platform independent if using the associated Docker/Singularity images.

Mac OS X or Linux CLI if run outside Docker/Singularity images.

Programming language: Bash, R 3.5, Perl, Python.

Other requirements: Python3.6 or higher, Java JDK, Chrome, Firefox, and Safari web browser.

License: MIT

Any restrictions to use by non-academics: None.

## Availability of data and materials

The dataset analyzed during the current study as a case study is publicly available at https://github.com/omicscodeathon/cdk1_crc/data. The data supporting the results reported in this manuscript is included within the article and its additional files. The generated progress reports are in HTML format and can be viewed using any preferred browser such as Chrome, Safari, Internet Explorer and Firefox. The Project repository which also includes the entire code and other requirements can be downloaded from https://github.com/omicscodeathon/cdk1_crc. The guidelines for implementing this tool and related updates, are available at: https://github.com/omicscodeathon/cdk1_crc/blob/master/README.md.

## Acknowledgements

The authors thank the National Institutes of Health (NIH) Office of Data Science Strategy (ODSS), the National Center for Biotechnology Information (NCBI) and the Genetics Society of America for their immense support before and during the October 2022 Omics codeathon organised in collaboration with the African Society for Bioinformatics and Computational Biology (ASBCB).

## Author Contributions

UCO conceived the original idea. UCO, OAE, PG, IA developed the bioinformatics pipeline. OAE, PG, EAI and IA performed the bioinformatic analysis of the case study data. UCO, OAE, CHO drafted the manuscript. OAE and IA intensively tested the pipeline and provided feedback. UCO, CHO, PWG, SO and HCA reviewed the manuscript and provided critical feedback and helped shape the final version of the manuscript. OIA’s role was in the administration and supervision of the bioinformatics analysis in the project. OIA also provided the resources to facilitate and complete the analysis and provided guidance. OIA did the editing and final review of the manuscript. All authors read and approved the final manuscript.

## Declarations

### Ethics approval and consent to participate

Not applicable.

### Consent for publication

Not applicable.

